# A novel plasmid-mediated PAC ß-lactamase (PAC-2) variant confers resistance to ceftazidime-avibactam in *Klebsiella quasipneumoniae* ST526, in Colombia

**DOI:** 10.1101/2025.05.14.654127

**Authors:** Sandra Yamile Saavedra, Efrain Montilla-Escudero, María Victoria Ovalle, Jeisson Alejandro Triana, Yeison Stid Torres, Nathalia Vargas-Flórez, María Alejandra Gutiérrez, Andrés Felipe Barrera, Carolina Duarte

## Abstract

Globally, the increasing reports of resistance to ceftazidime-avibactam (CZA) in Enterobacterales are concerning. In this paper we describe an isolate of *Klebsiella quasipneumoniae* (GMR-RA-2034.23) highly resistant to ceftazidime-avibactam, ceftolozane-tazobactam, ceftazidime and cefotaxime in which a new variant of the PAC ß-lactamase, designated PAC-2 by the NCBI, was identified. The *bla*PAC-2 gene is encoded in a non-conjugative plasmid type IncQ (9,452 bp). The GMR-RA-2034.23 isolate did not present mutations in the *OmpK*35 or *OmpK*36 porins, associated with resistance to CZA. This is the first description of PAC-type ß-lactamase in Colombia. Continuous monitoring of mechanisms of resistance to CZA is crucial to prevent its spread.

## Communication

Ceftazidime-avibactam (CZA) is a recently introduced ß-lactam/ß-lactamase inhibitor (BL/BLI) combination used to treat infections caused by carbapenemases-producing Enterobacterales (primarily KPC, OXA-48) and difficult-to-treat (DTR) resistance *Pseudomonas aeruginosa* (1, 2).

Reports of resistance to CZA have been increasing worldwide. They are associated with multiple mechanisms such as: (i) KPC variants with mutations affecting hotspots around the active site, mainly the Ω-loop (3, 4). (ii) ESBL-type ß-lactamases as: some variants of CTX-M with mutations in the Ω-loop (e.g., substitution P167S in CTX-M-14) (5), some variants of VEB such as VEB-25 (substitution K234R in VEB-1) (6), PER (7), co-existence of variants of GES (e.g., GES-26 and GES-19) (8). (iii) AmpC such as: PAC-1 (9), some variants of CMY (e.g., CMY-192) (10), mutations and hyperexpression of AmpC ß-lactamases (11) and (iv) a combination of multiple mechanisms, such as alterations in membrane permeability, due to porin loss or overexpression of efflux pump or expression of ß-lactamases (12, 13). PAC (*Pseudomonas aeruginosa* class C) is an unusual ß-lactamase, associated with resistance to CZA, identified in *Pseudomonas aeruginosa* and *Aeromonas* spp. Here, we realized the first report of PAC-2 in a clinical isolate of *Klebsiella quasipneumoniae*.

In December 2023, the National Reference Laboratory (NRL) of Colombia (Instituto Nacional de Salud “INS”), received a clinical isolate of *Klebsiella pneumoniae* (GMR-RA-2034.23) with resistance to CZA through the National Antimicrobial Resistance Surveillance program. The isolate was recovered from a bronchial secretion sample from a 76-year-old male patient, hospitalized in an Intensive Care Unit (ICU), the patient had received CZA as treatment for pneumonia caused by KPC-producing *Klebsiella pneumoniae*. GMR-RA-2034.23 was re-identified using matrix-assisted laser desorption ionization time-of-flight mass spectrometry (MALDI-TOF MS, Bruker). Antimicrobial susceptibility was confirmed by the disk diffusion assay and CZA susceptibility was determined by E-test (bioMérieux, Marcy l’Étoile, France). Susceptibility results were interpreted using the CLSI-M100 standard (14). The isolate was resistant to CZA (>256 μg/mL), ceftazidime, cefotaxime, cefepime, piperacillin-tazobactam, ceftolozane/tazobactam (C/T) and was susceptible to carbapenems (imipenem, meropenem and ertapenem), aztreonam, cefoxitin, gentamicin, ciprofloxacin, and amikacin. In GMR-RA-2034.23, the *bla*KPC, *bla*CTX-M, *bla*GES, *bla*PER, and *bla*CMY genes were assessed by PCR, yielding negative results for all ß-lactamases analyzed. Therefore, to determine the mechanism of resistance to CZA in the genome of the isolate, whole genome sequencing (WGS) was performed using Oxford Nanopore GidION technology (Oxford Nanopore Technologies, Oxford, UK).

For raw reads, the basecalling was conducted with Dorado (v0.7.1), de *novo* assemblies were performed using Unicycler (v0.5.0), the Coding sequence (CDS) annotation was carried out with Prokka (v1.14.6), the identification of resistance genes was performed using CARD Resistance Gene Identifier (RGI 6.0.1). Additional analyses were conducted with Kleborate (v2.4.1).

The WGS results indicate that GMR-2034.23 is a *Klebsiella quasipneumoniae* subspecies *quasipneumoniae* (genome size: 5,685,706 bp; GC content: 57.55%), belonging to ST526 being first report of this ST in Colombia. ST526, has been reported in clinical isolates from China and Singapore (15, 16) and non-clinical isolates from Brazil (17). GMR-RA-2034.23 harbors two ß-lactamases OKP-A-11 (OKP-like Species-specific chromosomal marker) and a new PAC allele (class C ß-lactamase), designated by NCBI as PAC-2 (PQ567105.1), for the first time reported globally; PAC-2 shares 99% identity with PAC-1 and differs by three substitutions in its amino acids (K44Q, E309A and N339K).

PAC-1 ß-lactamases have been associated with resistance to CZA and C/T, this ß-lactamase has been reported mainly in clinical isolates of *P. aeruginosa* from France (in patients repatriated from Mauritius and Afghanistan) (18), Nepal (19) and according to the NCBI pathogen isolates “PAC-1” database it has also been identified in countries such as Australia, Bangladesh, India, Israel, Spain, Singapore, Pakistan, Greece, United Kingdom, Kingdom of the Netherlands, and United States (https://www.ncbi.nlm.nih.gov/pathogens/isolates/#blaPAC-1%20 accessed 26 February 2025). PAC-1 has also been detected, in *Aeromonas caviae* and *Aeromonas enteropelogenes* recovered from a dog in Afghanistan (18) and a water reservoir in Singapore (20) respectively.

Literature reports indicate that in *P. aeruginosa* and *Aeromonas* spp., *bla*PAC-1 has been located on chromosome included in a group of conserved genes ISKpn9-PAC-1-2orfs-qacEΔ1-sul1 (18, 20) and only *bla*PAC-1 was found on the plasmid pGSH8M-1-1 (153.8 kb; AP019196.1) of an *A. caviae* isolate recovered from the effluent of an urban wastewater treatment plant in Japan (21). In this study *bla*PAC-2 was harbored on an IncQ-like plasmid with size 9,452 bp, the plasmid pKq-GMR-RA-2034.23 consists of the following structure: RepA (helicase), RepF (relaxase), MobA/RepB (fusion gene of relaxase–primase), MobC (protein required DNA cleavage), hypothetical protein and other proteins (Figure 1).

**Figure 1.**
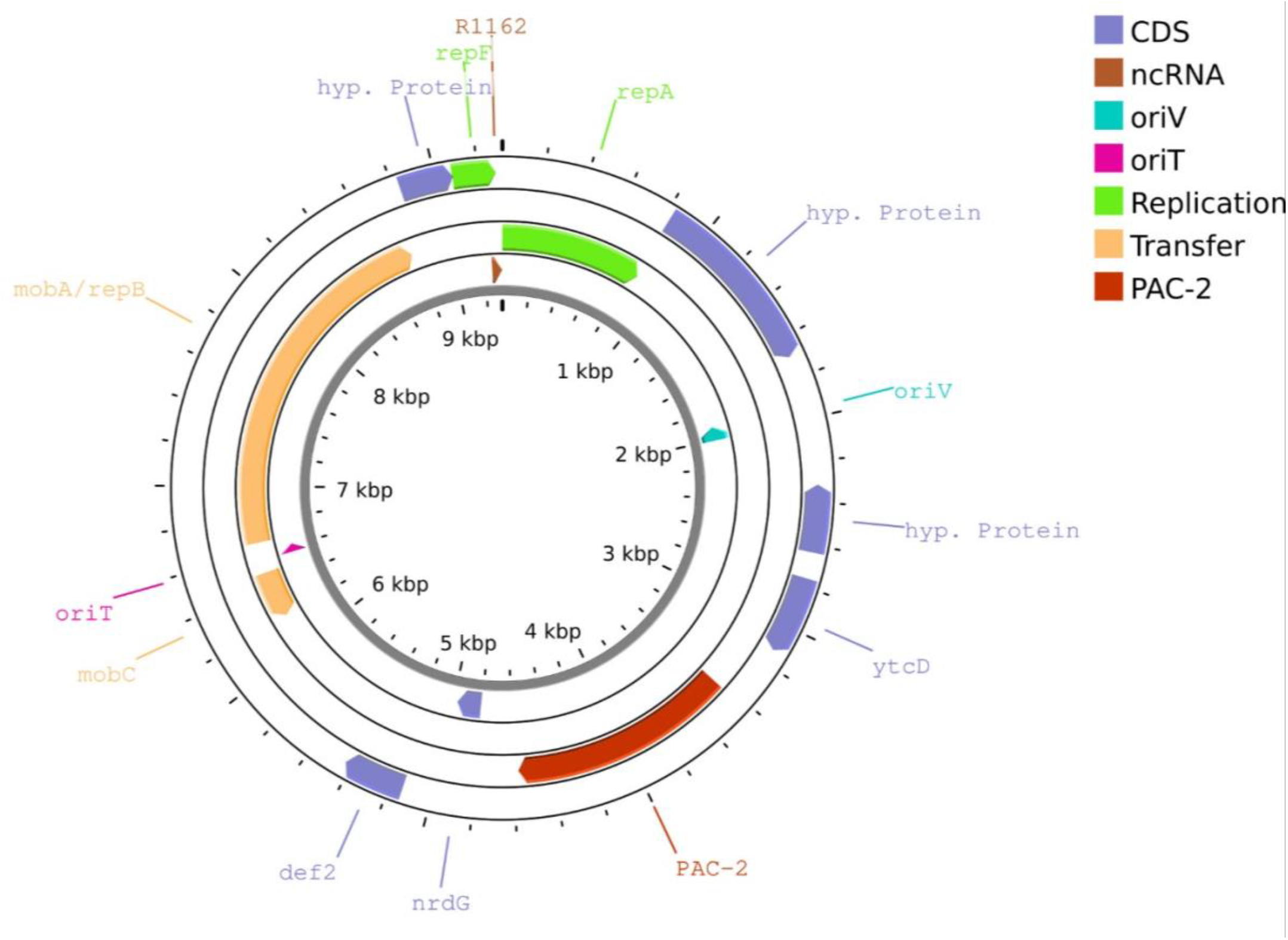
Schematic representation of plasmid incQ-like pKq-GMR-RA-2034.23 harboring PAC-2.

Conjugation experiments were unsuccessful, the IncQ plasmids are inherently non-self-conjugative but can be mobilized with the assistance of large conjugative plasmids. Due to their small size, high copy number, and broad host range replication, they have the potential to spread across diverse bacterial species (22). To our knowledge, this is the first global report of *bla*PAC-2, which is harbored in a plasmid. The plasmid pKq-GMR-RA-2034.23, show query coverage of 52% (identity of 99.18%) with the *Klebsiella pneumoniae* plasmid pCHE-A1 (KX244760) which carries *bla*GES-5 and aacA4 (23) (Figure 2) and a query coverage of 57% (identity of 96.40%) with the *Citrobacter werkmanii* plasmid pKPC-LB887 (MT569433) (24), (Figure 2), previously reported from South Africa and Brazil respectively.

**Figure 2.**
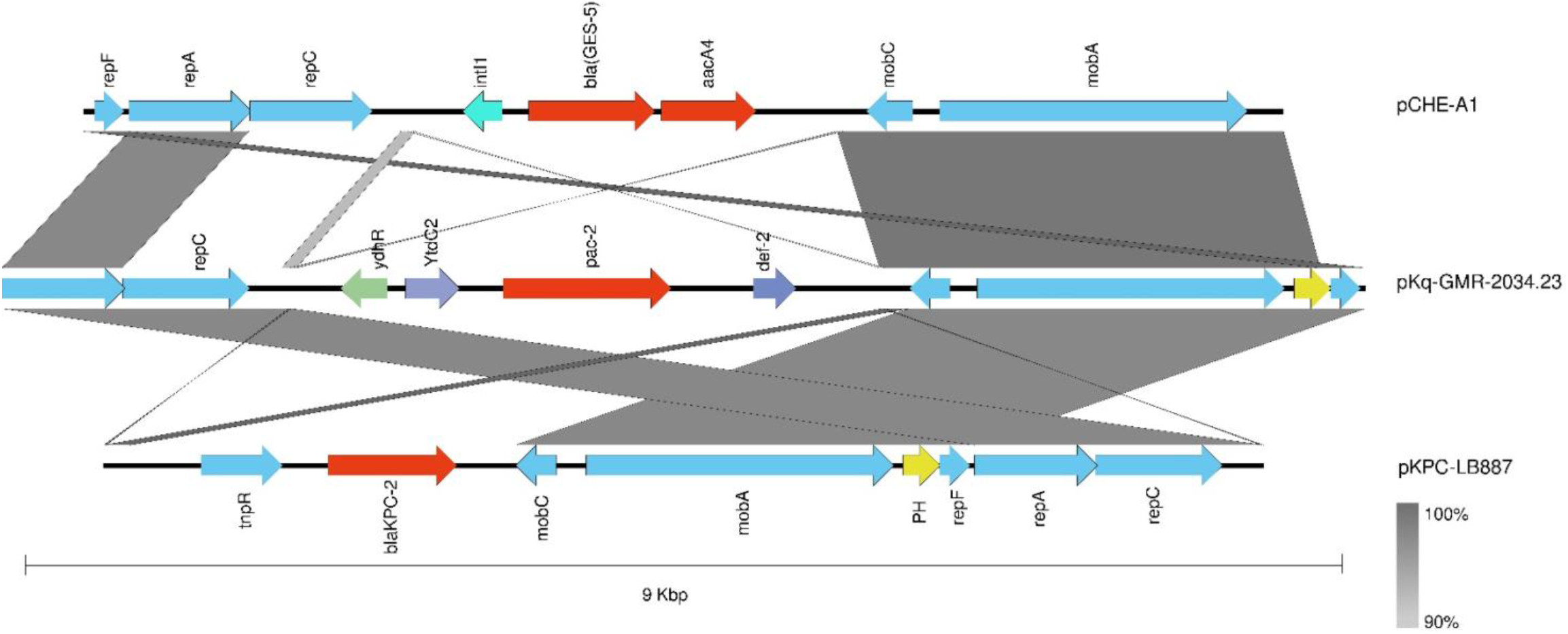
Schematic representation of comparative analysis of related plasmids. pKq-GMR-RA-2034.23 compared to plasmids pCHE-A1 (KX244760) and pKPC-LB887 (MT569433). Arrows indicate the direction of transcription. Red arrows represent the antimicrobial resistant gene. Light blue arrows replication genes, transfer genes and transposon-encoding genes. Yellow arrows Hypotetical proteins. Grey arrows and purple arrows represent other proteins.

β-lactamases associated with CZA resistance, such as PAC, CMY variants (e.g., CMY-192) (10), ESBL variants (CTX-M-14 “P167S”, PER, VEB-25) (5-7) and certain KPC variants (KPC-95, KPC-123, KPC-197) (4, 25) exhibit a cephalosporinase phenotypic profile and susceptibility to carbapenems, which represents a challenge in the laboratory for their phenotypic detection since they can only be identified using molecular tests. For β-lactamases such as KPC and CTX-M, detection is feasible using automated platforms or lateral flow immunochromatography tests; in contrast, β-lactamases such as VEB and PER are not available for analysis on automated platforms, while for PAC β-lactamases, there are no primers described in the literature and they have only been identified through WGS. Therefore, from the NRL we propose the primers PAC-F (5’-GTTTCTTTGGTCGTGGGCC-3’) and PAC-R (5’-CACTGGGCATGAAACACTCC-3’), for the performance of routine surveillance of PAC in isolates of CZA-resistant Enterobacterales.

Colombia is an endemic country for KPC and *Klebsiella pneumoniae* is the most frequently isolated microorganism in adult ICUs according to Whonet data from the NRL for 2023 (https://app.powerbi.com/view?r=eyJrIjoiNjRhYmRjYmItMGMwZS00NWEyLTgzNGMtZTMyNDRmYzBmMTZmIiwidCI6ImE2MmQ2YzdiLTlmNTktNDQ2OS05MzU5LTM1MzcxNDc1OTRiYiIsImMiOjR9&pageName=b07b0d6690c05e1000bd). In ICUs carbapenem resistance rates in *K. pneumoniae* within ICUs were 28.3% for imipenem (n=15,473), 25.9% for meropenem (n=18,800), and 27.5% for ertapenem (n=17,700) (data not shown). These findings underscore the critical role of CZA in the country as a treatment for infections caused by carbapenem-resistant Enterobacterales, such as *K. pneumoniae*.

In Colombia, the detection of different resistance mechanisms to CZA such as KPC variants (KPC-31 and KPC-197), reported since May 2024 (25) and PAC-2, threatens the use of CZA as the treatment of choice in infections by KPC-producing Enterobacterales, less than 5 years since its approval in Colombia. Given this concern, the NRL conducts continuous surveillance of CZA resistance and has published guidelines and recommendations (https://www.ins.gov.co/BibliotecaDigital/comunicado-tecnico-vigilancia-intensificada-por-laboratorio-de-resistencia-a-ceftazidimaavibactam-mediada-por-betalactamasas-en-enterobacterales-en-colombia.pdf) to enable laboratories to detect and report CZA resistance early. This allows timely preventive measures to be taken to contain the spread of these resistance mechanisms in the hospital setting, since combinations BL/BLI such as imipenem-relebactam, are in the process of being implemented in Colombia, while alternative treatment options such as cefiderocol are approved

## ACKNOWLEDGEMENTS

We acknowledge all public health laboratories from the national network of laboratories, which report to the National Reference Laboratory, Instituto Nacional de Salud “INS” of Colombia.

## REFERENCE

1. Papp-Wallace K. M. 2019. The latest advances in β-lactam/β-lactamase inhibitor combinations for the treatment of Gram-negative bacterial infections. Expert opinion on pharmacotherapy, 20: 2169–2184. 10.1080/14656566.2019.1660772

2. Tamma, P. D., Heil, E. L., Justo, J. A., Mathers, A. J., Satlin, M. J., Bonomo, R. A. 2024. Infectious Diseases Society of America 2024 Guidance on the Treatment of Antimicrobial-Resistant Gram-Negative Infections. Clinical infectious diseases: an official publication of the Infectious Diseases Society of America, ciae403. Advance online publication. 10.1093/cid/ciae403

3. Ding L, Shen S, Chen J, Tian Z, Shi Q, Han R, Guo Y, Hu F. 2023. Klebsiella pneumoniae carbapenemase variants: the new threat to global public health. Clin Microbiol Rev. 36:e0000823. 10.1128/cmr.00008-23

4. Hobson CA, Pierrat G, Tenaillon O, Bonacorsi S, Bercot B, Jaouen E, Jacquier H, Birgy A. 2022. Klebsiella pneumoniae carbapenemase variants resistant to ceftazidime-avibactam: an evolutionary overview. Antimicrob Agents Chemother 66:e0044722. 10.1128/aac.00447-22

5. Both A, Büttner H, Huang J, Perbandt M, Belmar Campos C, Christner M, Maurer FP, Kluge S, König C, Aepfelbacher M, Wichmann D, Rohde H. 2017. Emergence of ceftazidime/avibactam non-susceptibility in an MDR Klebsiella pneumoniae isolate. J Antimicrob Chemother. 72:2483–2488. 10.1093/jac/dkx179

6. Voulgari E, Kotsakis SD, Giannopoulou P, Perivolioti E, Tzouvelekis LS, Miriagou V. 2020. Detection in two hospitals of transferable ceftazidime-avibactam resistance in Klebsiella pneumoniae due to a novel VEB β-lactamase variant with a Lys234Arg substitution, Greece, 2019. Euro Surveill. 25:1900766. 10.2807/1560-7917.ES.2020.25.2.1900766

7. Poirel L, Ortiz de la Rosa JM, Sadek M, Nordmann P. 2022. Impact of Acquired Broad-Spectrum β-Lactamases on Susceptibility to Cefiderocol and Newly Developed β-Lactam/β-Lactamase Inhibitor Combinations in Escherichia coli and Pseudomonas aeruginosa. Antimicrob Agents Chemother. 66:e0003922. 10.1128/aac.00039-22

8. Khan A, Tran TT, Rios R, Hanson B, Shropshire WC, Sun Z, Diaz L, Dinh AQ, Wanger A, Ostrosky-Zeichner L, Palzkill T, Arias CA, Miller WR. 2019. Extensively Drug-Resistant Pseudomonas aeruginosa ST309 Harboring Tandem Guiana Extended Spectrum β-Lactamase Enzymes: A Newly Emerging Threat in the United States. Open Forum Infect Dis. 6:ofz273. 10.1093/ofid/ofz273

9. Bour M, Fournier D, Jové T, Pouzol A, Miltgen G, Janvier F, Jeannot K, Plésiat P. 2019. Acquisition of class C β-lactamase PAC-1 by ST664 strains of Pseudomonas aeruginosa. Antimicrob Agents Chemother. 2019 63:e01375–19. 10.1128/AAC.01375-19

10. Xu T, Wu W, Huang L, Liu B, Zhang Q, Song J, Liu J, Li B, Li Z, Zhou K. 2024. Novel plasmid-mediated CMY variant (CMY-192) conferring ceftazidime-avibactam resistance in multidrug-resistant Escherichia coli. Antimicrob Agents Chemother. 29:e0090624. 10.1128/AAC.01375-19

11. Xu T, Guo Y, Ji Y, Wang B, Zhou K. 2022. Epidemiology and mechanisms of Ceftazidime–Avibactam resistance in gram-negative bacteria. Engineering. 11:138–145. 10.1016/j.eng.2020.11.004

12. Chalhoub H, Sáenz Y, Nichols WW, Tulkens PM, Van Bambeke F. 2018. Loss of activity of ceftazidime-avibactam due to MexAB-OprM efflux and overproduction of AmpC cephalosporinase in Pseudomonas aeruginosa isolated from patients suffering from cystic fibrosis. Int J Antimicrob Agents. 52:697–701. 10.1016/j.ijantimicag.2018.07.027

13. Shi Q, Han R, Guo Y, Yang Y, Wu S, Ding L, Zhang R, Yin D, Hu F. 2022. Multiple Novel Ceftazidime-Avibactam-Resistant Variants of blaKPC-2-Positive Klebsiella pneumoniae in Two Patients. Microbiol Spectr. 10:e0171421. 10.1128/spectrum.01714-21

14. Clinical and Laboratory Standards Institute (CLSI). 2024. CLSI supplement M100. Performance standards for antimicrobial susceptibility testing. 34rd ed. Clinical and Laboratory Standards Institute, Wayne, PA.

15. Ma Z, Qian C, Yao Z, Tang M, Chen K, Zhao D, Hu P, Zhou T, Cao J. 2024. Coexistence of plasmid-mediated tmexCD2-toprJ2, blaIMP-4, and blaNDM-1 in Klebsiella quasipneumoniae. Microbiol Spectr. 12:e0387423. 10.1128/spectrum.03874-23

16. Octavia S, Kalisvar M, Venkatachalam I, Ng OT, Xu W, Sridatta PSR, Ong YF, Wang L, Chua A, Cheng B, Lin RTP, Teo JWP. 2019. Klebsiella pneumoniae and Klebsiella quasipneumoniae define the population structure of blaKPC-2 Klebsiella: a 5-year retrospective genomic study in Singapore. J Antimicrob Chemother. 74:3205–3210. 10.1093/jac/dkz332

17. Furlan JPR, da Silva Rosa R, Ramos MS, Dos Santos LDR, Savazzi EA, Stehling EG. 2024. Emergence of carbapenem-resistant Klebsiella pneumoniae species complex from agrifood systems: detection of ST6326 co-producing KPC-2 and NDM-1. J Sci Food Agric. 104:7347–7354. 10.1002/jsfa.13555

18. Bour M, Fournier D, Jové T, Pouzol A, Miltgen G, Janvier F, Jeannot K, Plésiat P. 2019. Acquisition of class C β-lactamase PAC-1 by ST664 strains of Pseudomonas aeruginosa. Antimicrob Agents Chemother. 63:e01375–19. 10.1128/AAC.01375-19

19. Tada T, Shimada K, Satou K, Hirano T, Pokhrel BM, Sherchand JB, Kirikae T. 2017. Pseudomonas aeruginosa Clinical Isolates in Nepal Coproducing Metallo-β-Lactamases and 16S rRNA Methyltransferases. Antimicrob Agents Chemother. 61:e00694–17. 10.1128/AAC.00694-17

20. Zhong Y, Guo S, Thong S, Schlundt J, Kwa AL. 2023. First report of environmental blaPAC-1-carrying Aeromonas enteropelogenes. Microbiol Spectr. 11:e0139123. 10.1128/spectrum.01391-23

21. Sekizuka T, Inamine Y, Segawa T, Hashino M, Yatsu K, Kuroda M. 2019. Potential KPC-2 carbapenemase reservoir of environmental Aeromonas hydrophila and Aeromonas caviae isolates from the effluent of an urban wastewater treatment plant in Japan. Environ Microbiol Rep. 11:589–597. 10.1111/1758-2229.12772

22. Loftie-Eaton W, Rawlings DE. 2012. Diversity, biology and evolution of IncQ-family plasmids. Plasmid. 67:15–34. 10.1016/j.plasmid.2011.10.001

23. Pedersen T, Sekyere JO, Govinden U, Moodley K, Sivertsen A, Samuelsen Ø, Essack SY, Sundsfjord A. 2018. Spread of Plasmid-Encoded NDM-1 and GES-5 Carbapenemases among Extensively Drug-Resistant and Pandrug-Resistant Clinical Enterobacteriaceae in Durban, South Africa. Antimicrob Agents Chemother. 62:e02178–17. 10.1128/AAC.02178-17

24. Campana EH, Kraychete GB, Montezzi LF, Xavier DE, Picão RC. 2022. Description of a new non-Tn4401 element (NTEKPC-IIe) harboured on IncQ plasmid in Citrobacter werkmanii from recreational coastal water. J Glob Antimicrob Resist. 29:207–211. 10.1016/j.jgar.2023.07.015

25. De la Cadena E, Mojica MF, Rojas LJ, Castro BE, García-Betancur JC, Marshall SH, Restrepo N, Castro-Caro NP, Fonseca-Carrillo M, Pallares C, Bonomo RA, Villegas MV. 2024. First report of KPC variants conferring ceftazidime-avibactam resistance in Colombia: introducing KPC-197. Microbiol Spectr. 12:e0410523. 10.1128/spectrum.04105-23

